# KYLO-0603, a novel liver-targeting, thyroid hormone receptor-β agonist for the treatment of MASH

**DOI:** 10.1101/2025.02.09.637336

**Authors:** Xueqin lu, Shengjun Wang, Yanchun Du, Bixian Xie, Qingyan Chen, Jinzhen Lin, Bailing Chen, Kunyuan Cui

## Abstract

Metabolic dysfunction–associated steatohepatitis (MASH) is a progressive liver disease associated with liver-related complications and death. Kylo-0603 is a novel agonist for the thyroid hormone receptor β (THR-β) that has been developed by merging the structures of three acetylgalactosamine (GalNAc)-modified ASPGR ligands with a triiodothyronine (T3) analog. This unique design allows for both THR-β activation and targeted delivery to hepatocytes, significantly reducing the risk of adversed effects related to increased systemic thyroid hormone-like effects. Additionally, it effectively lowers serum cholesterol by as much as 69.2% and low-density lipoprotein cholesterol (LDL-C) levels by up to 88.2% in the MASH mouse model. Meanwhile, Kylo-0603 was shown to reduce steatosis by up to 1.3 points (P < 0.001), inflammation by 1.8 points (P < 0.0001), and ballooning by 0.8 points (P < 0.01). The non-alcoholic steatohepatitis (NASH) activity score (NAS) decreased by up to 3.7 points (P < 0.0001), and the fibrosis score dropped by 0.6 points (P < 0.05). These findings suggest that Kylo-0603 is effective in enhancing liver tissue NASH status and inhibiting fibrosis progression. In summary, Kylo-0603, as a highly both tissue and target selective and low-toxicity THR-β agonist, shows significant promise for treating MASH and is likely to emerge as a new therapeutic option for patients with this condition.

## 1. Introduction

The liver is an essential organ closely related to lipid metabolism and is responsible for both the regulation of lipid homeostasis and energy utilization [1, 2]. Metabolic dysfunction-associated steatotic liver disease (MASLD), formerly known as nonalcoholic fatty liver disease (NASLD), can be diagnosed in adults with hepatic steatosis detected by imaging techniques, blood biomarkers, or liver histology, when overweight or obese, or in the presence of T2DM or at least two metabolic risk abnormalities [3]. Currently, MASLD is the most common liver disease worldwide, affecting 30% of the adult population. According to regional prevalence data, the highest rate of MASLD is found in the Middle East (32%) and South America (30%), and the lowest is found in Africa (13%), but prevalence rates are even higher in specific subpopulations such as severely obese patients (90%) and patients with type 2 diabetes (76%). Although liver steatosis is usually benign, it can progress to MASH, previously termed nonalcoholic steatohepatitis (NASH) [4, 5]. MASH is an important sign of disease progression, and, if not controlled, can progress to cirrhosis and even liver cancer [6, 7, 8, 9, 10]. In contrast to simple steatosis where the primary histological feature is lipid accumulation in liver cells, MASH is characterized by additional liver inflammation with or without fibrosis, Chronic steatosis drives the disease progression toward MASH, which is characterized by inflammation and ballooning or hepatocyte damage [11, 12, 13]. In recent decades, many researchers and pharmaceutical companies have made great efforts to develop drugs for MASLD and MASH in an attempt to inhibit the MASH process. Despite extensive research into the pathogenesis and drug development of MASH, only one drug that targets the treatment of MASH was approved by the FDA (U.S. Food and Drug Administration) in 2024 [7, 14, 15, 16].

Thyroid hormones (THs) are considered to be important signaling molecules for maintaining normal body metabolism and play critical roles in differentiation, growth, and metabolism [17, 18]. They exert their physiological effects by binding to specific nuclear receptors, THR-α and THR-β, which are differentially expressed and control tissue/cell-specific thyroxine activity [19-20]. In contrast to the THR-α receptor, which is predominantly expressed in the heart, the THR-β isoform is primarily expressed in the liver and exerts significant effects on lipid metabolism [22, 23]. Specifically, THR- β has been demonstrated to reduce LDL-C levels, decrease generalized obesity and body weight, and is capable of reducing lipid content by increasing the rate of lipid metabolism in the liver [24, 25]. Moreover, Perra et al. demonstrated that THR agonists have the potential to inhibit or reverse hepatic steatosis [26]. This significant discovery highlights the remarkable therapeutic potential of THR agonists in the treatment of MASLD. Several THR-β agonists have been studied, including Resmetirom, which has been approved for marketing by the FDA, but none of these compounds are liver-specific [20, 27-28].

The hepatic asialoglycoprotein receptor (ASGPR) is a highly abundant, rapidly internalized receptor with many cellular lectins selectively distributed on the surface of hepatocytes [29-31]. Modification of drug formulations with ASGPR ligands can significantly improve drug selectivity and permeability to hepatocytes[32-34]. ASGPR promotes the uptake and clearance of circulating glycoproteins with exposed terminal galactose and n-acetylgalactosamine, amino sugar derivatives of galactose and glycans, via lattice protein-mediated endocytosis [35-37]. Thus, a consensus has been reached that triantennary GalNAc with a mutual distance of ∼20 Å exhibits the highest affinity with ASGPR, which has been widely used for liver-targeted delivery of various payloads [38,39].

The present study was undertaken to mitigate the adverse effects of therapeutic agents for MASLD or MASH and to enhance the accumulation of THR-β agonists in the liver while maintaining the regulatory role of thyroxine on intrahepatic lipid metabolism. To this end, a series of innovative compounds incorporating the liver-targeting ligand Gal-NAc with thyroxine receptor agonists were designed and synthesized (see our patent). Following a period of careful screening, Kylo-0603 was identified as the optimal candidate (Figure 1). The data from the study demonstrated that, following an eight-week treatment period, Kylo-0603 resulted in a significant improvement in MASH and/or fibrosis in the MASH mouse model (F2∼3) induced with a high-fat diet (HFD) in conjunction with carbon tetrachloride (CCl_4_) induction while simultaneously reducing LDL-C and body weight. This provides an additional therapeutic advantage for patients with MASH associated with hyperlipidemia or obesity.

**Figure 1.**
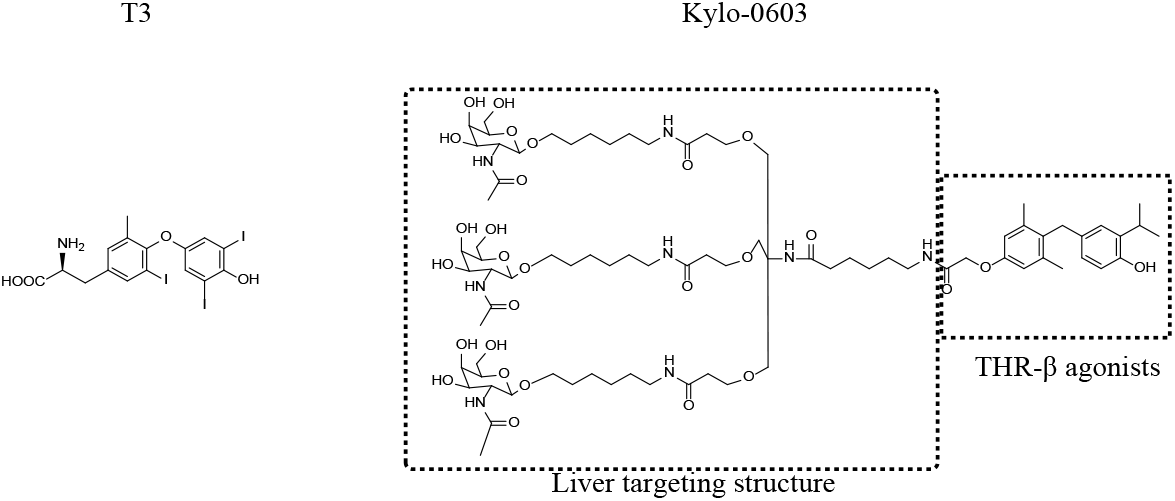
The structures of T3 and Kylo-0603

## 2. Materials and methods

### 2.1 Chemistry

All reactions were run under an inert atmosphere (Ar) with flame-dried glassware using standard techniques for manipulating air-sensitive compounds. All the solvents were dried and purified before use by standard procedures. Commercial reagents were used as supplied or purified by standard techniques where necessary. Column chromatography was performed using 200-300 mesh silica with the proper solvent system according to TLC analysis and UV light to visualize the reaction components. Unless otherwise noted, nuclear magnetic resonance (NMR) spectra were recorded on a 400 MHz spectrometer. NMR data were reported as follows: chemical shift, multiplicity (s = singlet, d = doublet, dd = doublet-doublet, t = triplet, q = quartet, qt= quartet of triplets, m = multiplet), coupling constant in Hz and integration. Chemical shifts of 1H NMR spectra were recorded in parts per million (ppm) on the δ scale from an internal standard of residual CDCl_3_ (7.26 ppm), and CD_3_CN (1.94 ppm). Chemical shifts for ^13^C NMR spectra were recorded in parts per million from tetramethylsilane using the central peak of CDCl_3_ (77.16 ppm), and CD_3_CN (1.32, 118.26 ppm) as the internal standard. HR-MS data were obtained using ESI ionization with 100,000 (FWHM) maximum resolution.

### 2.2 THR-β selectivity assessment of Kylo-0603

In conducting experiments to ascertain the agonist effects of compounds on human THR-α or THR-β, the following equipment and reagents were utilized: The experimental apparatus comprised an Envision instrument (provided by PerkinElmer, model Envision 2014) and an Echo 500 instrument from Labcyte, USA. The experimental reagents used were the LanthaScreen™ TR-FRET THRα/THRβ Coactivator Detection Kit (model PV4687) from Thermo Fisher Scientific. The procedure was as follows: initially, gradient dilutions of the drugs Kylo-0603 and T3 were conducted on the Echo 500 instrument. The dilution was conducted via a 10-point, 3-fold serial dilution, with the initial concentration of Kylo-0603 ranging from 400 μM to 40 μM and that of T3 ranging from 40 nM to 2 nM. Subsequently, 100 nL of the diluted compounds were transferred to a 384-well experimental plate (384 Optiplate), with one compound well designated for each concentration. The experimental procedure was conducted by the instructions provided in the LanthaScreen™ TR-FRET TRα/TRβ Coactivator Assay Kit. To ensure the accuracy and reliability of the data, each experiment was repeated three times. EC50 values were determined by using GraphPad Prism 8.5 (the Dose-response-Stimulation-log[effect] vs. response--Variable slope). Finally, to assess the selectivity of the compounds, we calculated the selectivity multiplicity. Fold of selectivity was calculated by (compound EC50 on THR-α/T3 EC50 on THR-α) / (compound EC50 on THR-β/T3 EC50 on THR-β)

### 2.3 Animal study

this experimental protocol and any changes to the experimental protocol involving the use and care of animals were reviewed and approved by the Institutional Animal Care and Utilization Committee (IACUC) committee before implementation. During the experiments, animal welfare and experimental procedures followed AAALAC animal welfare requirements. The animals were maintained on a 12/12 h light/dark cycle with free access to water and food. All animals come from GemPharmatech Co., Ltd.

#### 2.3.1 Experimental observation and data collection of HFD-induced obese mice

Experiments were performed on 112 (n = 16/group) male C57BL/6J mice (5 weeks). After 3 days of acclimatization, the animals were randomly divided according to body weight into a control group (Chow Diet) and a HFD group (HFD, 60% calories as fat, Cat# D12492, Research Diet, Jiangsu Xietong Pharmaceutical Bioengineering Co., Ltd.). The mice were provided with the corresponding diets for 16 weeks. The mice were fed the corresponding diets for 16 weeks (with bedding shavings and a high-fat diet changed twice a week), during which time their weights were recorded once a week. In the meantime, Kylo-0603 (0.1 mg/kg to 10 mg/kg) or vehicle was orally administered to HFD-induced obese mice once daily. A group of mice fed with a normal diet was set as a normal control and was gavaged with a vehicle once every day. At the end of the experiment, the mice were fasted for 6 hours and then sacrificed. Blood samples were collected and then centrifuged to obtain the serum for the detection of cholesterol, triglyceride, and low-density lipoprotein cholesterol. The heart and liver tissues were excised from each group of mice and weighed and photographed. The hearts were frozen in liquid nitrogen at -80°C. The livers and hearts of the mice were rapidly removed and washed with cold saline. Half of the left lobe was fixed in 4% neutral paraformaldehyde, and the remaining livers and hearts were snap-frozen and stored at -80°C for further analysis. The concentrations of lipids and liver enzymes were determined by separating serum from orbital or cardiac blood and measuring triglycerides (TG), cholesterol (Chol), Low-density lipoprotein cholesterol (LDL-C), alanine transaminase (ALT), and aspartate transaminase (AST) levels using a Hitachi 7020 automated hematology laboratory and a Wako kit.

#### 2.3.2 Experimental observation and data collection of MASH mouse model (F2∼3) induced with a high-fat diet (HFD) in conjunction with carbon tetrachloride (CCl_4_) induction

A total of 80 5-week-old C57BL/6J male mice were randomly assigned to two groups based on their body weight: a control group (n = 10) and a high-fat diet group (HFD, D12492, n = 70). During the eight-week feeding period, the mice in the two groups were provided with the corresponding diets, including shavings bedding, and their weights were recorded once a week. The high-fat diet was changed twice weekly. Weighing the weight of each group of mice at a given time. The 60 mice with the highest body weights in the 60% high-fat diet group were subsequently selected and randomly divided again into six groups of 10 mice each based on the base of body weight. The control group was provided with a control diet supplemented with the solvent, whereas the model group was given a high-fat diet with the same supplement. The remaining four groups were provided a high-fat diet containing the solvent and a range of doses of Kylo-0603, namely, 0.1 mg/kg, 0.3 mg/kg, 1 mg/kg, 3 mg/kg, and 10 mg/kg. The control group was administered intraperitoneal injections of 0.2 ml/kg CCl_4_ twice a week for 8 weeks, during which time the mice were fed the same diets as before the grouping, and were weighed once a week. The mice were weighed weekly, and the dosage for each week was determined based on their body weight at the commencement of that week. On the day following the final administration of the drug (after a period of fasting of between 5 and 6 hours), the mice in each group were euthanized, whole blood was collected from either the orbits or the heart, and the serum was separated and stored at -80°C. The heart and liver tissues of the mice in each group were weighed and photographed. The heart tissues were then quickly frozen in liquid nitrogen and stored in a refrigerator at -80°C, and the middle lobe of the liver was divided into four parts. Two of these samples were quickly frozen in liquid nitrogen and stored at -80°C (for RNA sequencing). The remaining two portions were fixed in a 4% Paraformaldehyde solution and subsequently dehydrated for paraffin embedding and OCT embedding. The concentrations of lipids and liver enzymes were determined by separating serum from orbital or cardiac blood and TG, Chol, LDL-C, ALT, and AST levels via a Hitachi 7020 automated hematology laboratory and a Wako kit. The sera were separated from the blood collected via cardiac puncture in each group via Cloud-Clone products and tested according to the instructions provided with the thyroid stimulating hormone (TSH, also known as thyrotropin), triiodothyronine (T3), thyroxine (T4), free T4 (fT4) and free T3 (fT3) enzyme immunoassay kits. The liver tissue was thoroughly homogenized in isopropanol (1:9 tissue to isopropanol by mass/volume) on ice, left overnight at 4°C, and centrifuged to remove the supernatant. The resulting solution was analyzed for TG and Chol using a Hitachi 7020 automated hematochemistry system.

#### 2.3.3 Hematoxylin and eosin (H&E)

The dewaxed slides were placed in the alcohol-free hematoxylin staining solution for a period of 3-5 minutes. They were then rinsed thoroughly with flowing tap water. The slides were differentiated and returned to blue for several seconds, before being dehydrated sequentially with 50%, 70%, 85%, and 95% ethanol. They were then subjected to eosin staining for a period of 5-10 minutes. The slides were then dehydrated sequentially with 95%, 100%, and 100% ethanol before being cleared with xylene. Finally, the slides were sealed with neutral gum, dried, and imaged.

#### 2.3.4 Oil Red O Staining

The frozen sections were immersed in 60% isopropyl alcohol for 3–5 minutes, followed by immersion in Oil Red O staining solution for 3–5 minutes. The samples were then washed with pure water.

Following a wash with pure water, the nuclei were restained with hematoxylin, after which the sections were sealed with 50% glycerol, dried and photographed for imaging.

#### 2.3.5 Sirius Red Staining

The dewaxed slides were rinsed in running water, stained with Sirius Red for 8-10 minutes, washed slightly in anhydrous ethanol, dehydrated, made transparent by gradient ethanol and xylene, and finally sealed with neutral gum, dried and photographed for imaging.

#### 2.3.6 Statistical analysis

The results of the experiments, including the body weights of the animals in each group, are expressed as the means ± standard deviations (means ± SD). One-way analysis of variance (ANOVA) was employed to determine whether there were any significant differences between the various treatment groups and the control group. The data were analyzed via GraphPad Prism 8.5, and a p-value of less than 0.05 was considered statistically significant.

#### 2.3.7 SHG/TPEF microscopy and imaging procedure

Scanning was conducted on a Genesis® machine, a stain-free TPE/SHG microscopic imaging device manufactured in-house by HistoIndex®, Singapore. The region of interest (ROI) is divided into subregions of equal size, termed “blocks,” with standard dimensions of 200 μm × 200 μm. Each block is imaged individually and in sequence via raster scan mode with a 20x objective. Each block has a resolution of 512 × 512 pixels, or 0.4 μm per pixel. Software developed by HistoIndex and Chipmap for image analysis identifies the different lobular regions, the central vein, the confluent region, and the perisinus region. Fibrosis and steatosis in these regions were subsequently measured and quantified in the SHG and TPE images.

#### 2.3.8 PK studies

The selected compounds were orally administered to ICR rats (male, n = 3 per time point) separately. Blood samples were collected before and at 0.25, 0.5, 1, 2, 4, 8, and 24 h after oral administration and then centrifuged at 4500 rpm for 10 min at 4°C to obtain the serum. Liver samples were collected before and at 1, 4, 8, and 24 h after oral administration. The concentrations of the compounds in the serum or liver were determined via liquid chromatography/tandem mass spectrometry (LC−MS/MS).

#### 2.3.8 Gene expression analysis

Transcriptome sequencing was performed on the Illumina sequencing platform, and the library sequencing and biotechnological analysis were performed by Beijing Novozymes Technology Co.

## 3. Results

### 3.1 Design and synthesis of Kylo-0603

This study is focused on the design and synthesis of an innovative liver-targeting compound, Kylo-0603, which combines the highly liver-targeting properties of GalNAc with those of a thyroid hormone receptor agonist. Specifically, Kylo-0603 was prepared by chemically coupling Compound B with Compound A (a GalNAc derivative, the synthesis of which is shown in the supporting document). The synthesis of Kylo-0603 is shown in Figure 2. The specific experimental procedure was to sequentially add DMF (3.0 mL), Compound B (15 mg), TBTU (8.47 mg), and DIPEA (20.2 mg) to the reaction vessel and react for 6 hours. Compound A (47 mg) was then rapidly added and stirred for 2 hours at room temperature. The reaction was detected by high-performance liquid chromatography (HPLC), and the reaction was completed and terminated. The reaction mixture was prepared with a 1.0 mol/L ammonia solution in an ice bath to ensure that the pH of the reaction mixture was 8-10. The ice bath was removed, and the mixture was stirred at room temperature for half an hour while HPLC detection was performed. At the end of the reaction, the pH was adjusted to 7.0 with glacial acetic acid, and the mixture was then concentrated. The concentrated residue was dissolved in 35% acetonitrile/water, filtered and lyophilized to yield 29.47 mg of the target compound. ^1^H NMR (400 MHz, DMSO-d_6_) δ 8.97 (s, 1H), 8.01 (d, J= 5.2 Hz, 1H), 7.82 (t, J= 5.2 Hz, 3H), 7.65 (d, J= 8.8 Hz, 3H), 7.00 (s, 1H), 6.82 (s, 1H), 6.65 (s, 2H), 6.61 (d, J= 8.0 Hz, 1H), 6.45 (d, J= 7.6 Hz, 1H), 4.61 (t, J= 5.2 Hz, 3H), 4.57 (d, J= 4.8 Hz, 3H), 4.48 (d, J= 2.8 Hz, 3H), 4.40 (s, 2H), 4.24 (d, J= 8.0 Hz, 3H), 3.79 (s, 2H), 3.73 (m, 3H), 3.69 (m, 3H), 3.53 (m, 24H), 3.34 (m, 6H), 3.12 (m, 3H), 3.03(d, J= 5.2 Hz, 6H), 2.29 (t, J= 5.6 Hz, 6H), 2.15 (s, 6H), 2.07 (t, J= 7.2 Hz, 3H), 1.82 (s, 9H), 1.24-1.43 (m, 30H), 1.09 (d, J= 6.8 Hz, 6H). ^13^C NMR (100 MHz, DMSO-d6)) 172.98, 170.57, 170.09, 168.23, 155.91, 152.70, 138.12, 134.35, 130.92, 130.28, 125.88,125.35, 115.28, 114.52, 101.84, 75.65, 71.98, 68.79, 68.54, 68.05, 67.83, 67.37, 60.99, 59.98, 52.64, 38.99, 38.69, 36.46, 36.34, 33.58, 29.63, 29.53, 29.36, 26.91, 26.69, 26.40, 25.66, 25.50, 23.47, 22.93, 20.51; HR-MS (ESI) m/z calcd for C_81_H_134_N_8_O_28_ [M + H] ^+^: 1667.9380, found: 1667.9382.

**Figure 2.**
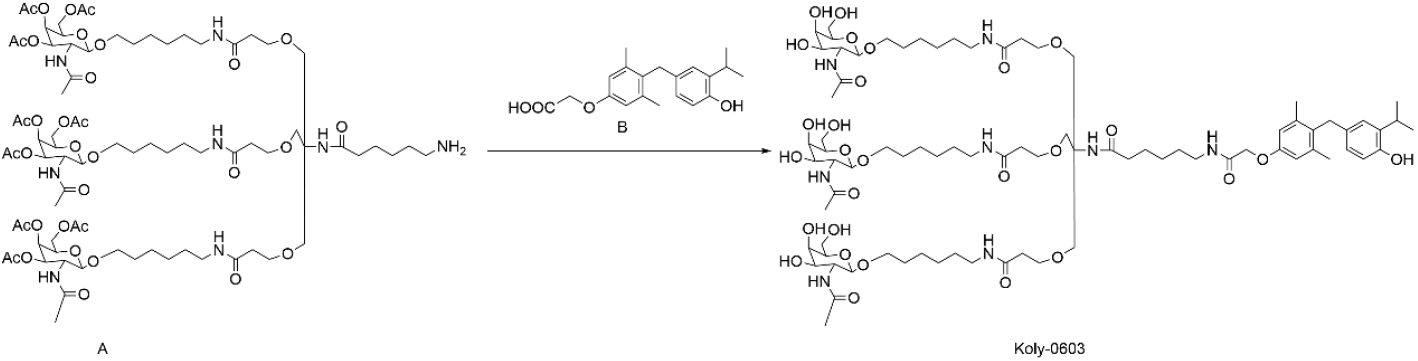
Synthesis process of Kylo-0603

### 3.2 Kylo-0603 drug stability determination and in vitro receptor binding assay

We first evaluated the metabolic stability of Kylo-0603 in the plasma of CD-1 mice, Sprague–Dawley (SD) rats, beagle dogs, cynomolgus monkeys, and humans. The plasma from the five species was mixed with Kylo-0603 (2 μM) and incubated at 37°C in a thermostatic water tank. The plasma concentrations of Kylo-0603 were monitored via LC-MS/MS at 0, 10, 30, 60, and 120 min. The half-lives (T1/2) of Kylo-0603 in the plasma of CD-1 mice, SD rats, beagle dogs, cynomolgus monkeys and humans were determined to be >372.7, >372.7, 355.8, >372.7 and >372.7 min, respectively. Kylo-0603 was stable in the plasma of CD-1 mice, SD rats, cynomolgus monkeys and humans and unstable in beagle dog plasma.

### 3.3 THR-β selectivity assessment of Kylo-0603

Kylo-0603 is a new compound based on a structural modification of the thyroid hormone T3 that incorporates GalNAc and a T3-like thyroid hormone structure, a design that gives Kylo-0603 the dual properties of a liv-er-targeting and THR-β receptor agonist. To assess the binding selectivity of Kylo-0603 for THR-α and THR-β receptors and compare it with that of T3, THR-α and THR-β receptor coactivator experiments were performed. The experimental results revealed that Kylo-0603 bound THR-α and THR-β with EC50 values of 124.1±13.92 nM and 31.07±4.42 nM, respectively, whereas T3 bound both receptors with EC50 values of 0.01±0.002 nM and 0.02±0.001 nM, respectively. After normalization for the relative selectivity of T3 binding to the receptors, the results revealed that the relative selectivity of Kylo-0603 for THR-β was 8.2-fold greater than that for THR-α (Figure 3A). Although Kylo-0603 presented a lower affinity ratio for THRα than for THR-β compared with the established compound GC-1, it nevertheless surpassed previous THRβ agonists in terms of hepatic targeting efficacy. To evaluate the pharmacokinetic properties of the novel compound Kylo-0603, a single dose of 7.03 mg/kg was administered to male Sprague-Dawley (SD) rats maintained under fasting conditions. The plasma and hepatic drug concentrations were accurately analyzed via high-performance liquid chromatography coupled with tandem mass spectrometry (HPLC-MS/MS). The results revealed a low plasma exposure level of Kylo-0603 (12.8 pg/ml at 0.25 h). In contrast, a notable accumulation of the drug was observed in the liver, where the concentration reached an exceptionally high level of 66167 pg/g at 0.25 hours post-dosing. This finding demonstrates that the drug has specific targeting properties in the liver (Figure 3C). Nevertheless, the ratio of tissue to plasma concentration was comparable for T3 (Ana Cachefo, 2001; Villicev et al., 2007).

**Figure 3.**
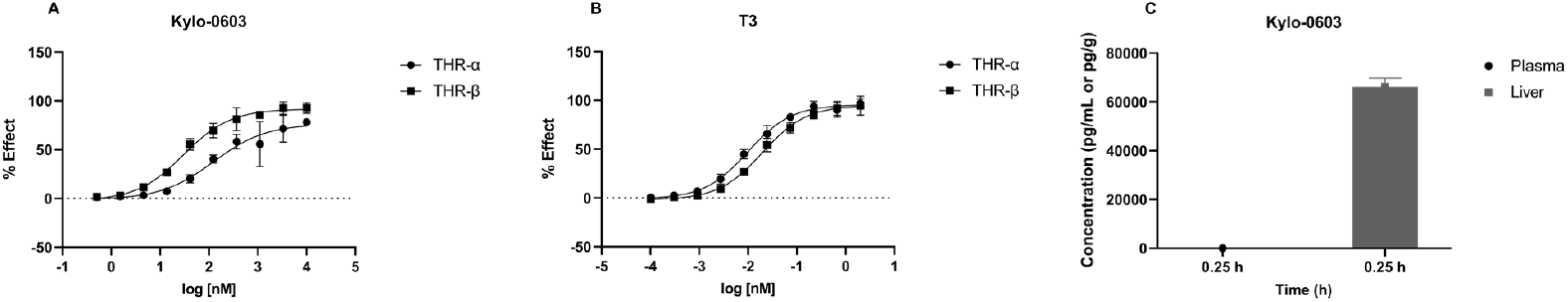
EC50 ratios of Kylo-0603 for the THR-α and THR-β receptors (A). The EC50 ratios of T3 to THR-α and THR-β receptors (B). Plasma and Liver Concentrations of Kylo-0603 after PO1 Dosing in Rats (C)

### 3.4 Effects of Kylo-0603 on body weight and fat content in HFD-induced obese mice

Compared with those in the normal diet control group (chow), the high-fat diet (HFD)-induced obesity model mice in this study presented significant increases in body weight, liver weight, body fat percentage, and body lean mass percentage. During the experimental period, food consumption did not significantly differ between the vehicle and experimental groups. After a single daily dose of Kylo-0603 for 10 weeks in conjunction with HFD feeding, the body weights of the mice in the 0.1, 0.3, 1, 3, and 10 mg/kg Kylo-0603 groups were 0.8%, 4.8%, 3.8%, 13.1%, and 15.8% lower than those of the mice in the HFD group, respectively (Figure 4A). The liver weight and body weight of the mice significantly decreased in all the dose groups, but the effect of various doses of Kylo-0603 on the heart weight of the mice was not significant (Figure 4B and Figure 4C). An interesting dose-response relationship was observed with the drug intervention: the 10 mg/kg dose significantly reduced the body weight of the mice, resulting in a significant increase in the heart weight/body weight ratio at this dose (Figure 4E). However, the liver weight decreased with increasing concentrations of the drug administered, so the liver weight/body weight ratio essentially remained unchanged at the low dose and increased only at the highest dose of 10 mg/kg (Figure 4D). Body length and bone density were not affected by Kylo-0603 (Figure s2).

**Figure 4.**
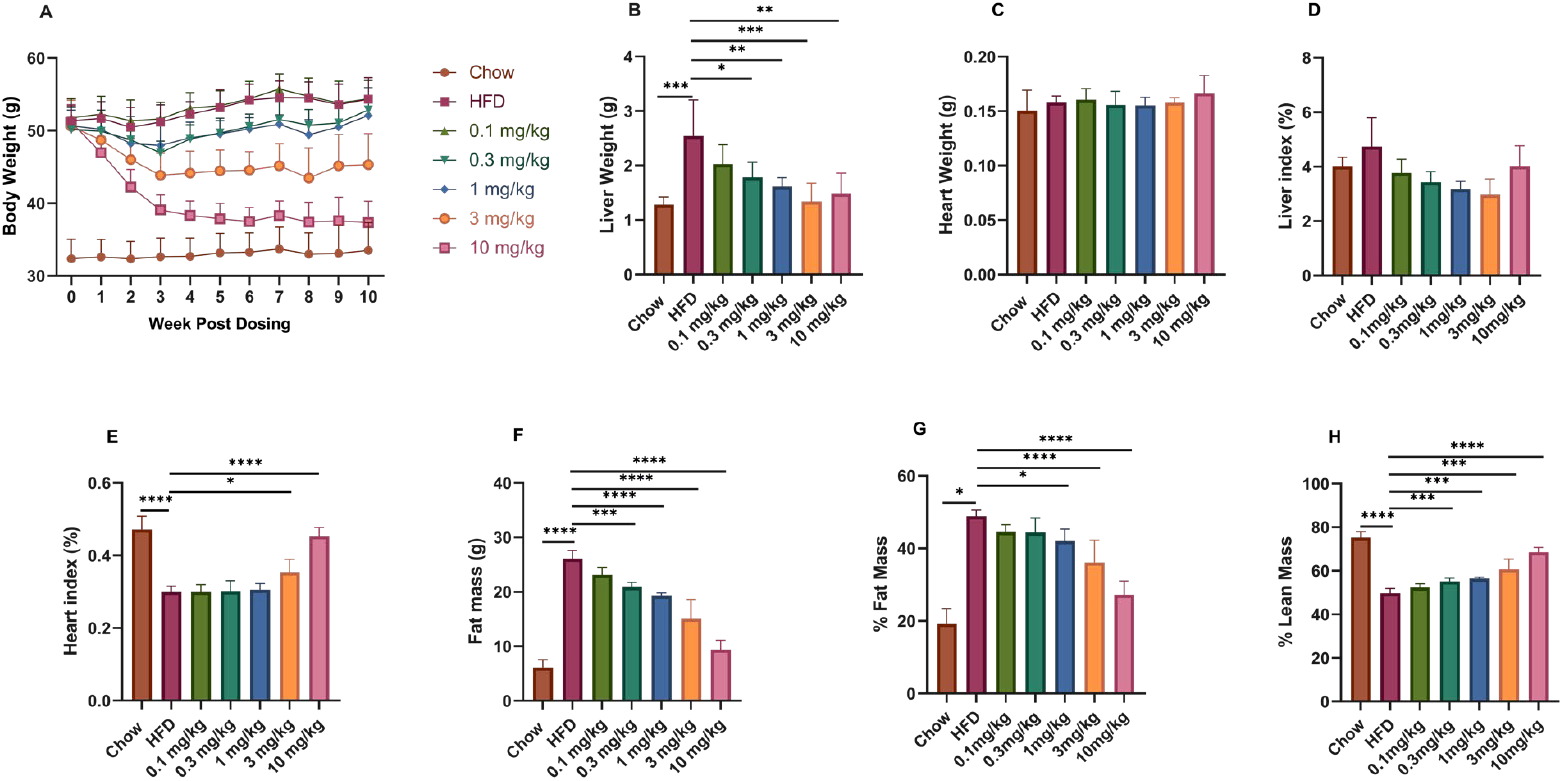
Effects of Kylo-0603 on body weight, organ weight, and body fat content. Male 5-week-old C57BL/6J mice were fed either a normal diet or a high-fat diet (HFD; 60 kcal% fat) and maintained for 16 weeks. The animals were subsequently administered a single daily oral dose of Kylo-0603 for 10 weeks in conjunction with HFD feeding. Body weights of the mice after drug intervention (A). Liver and heart weights of the mice after 10 weeks of drug intervention (B and C). Relative fat mass% (E) and lean mass% (F) after 8 weeks of Kylo-0603 treatment. Data are shown as the mean±SD, n=5; * P< 0.05, **P< 0.01, ***P< 0.001, and ****P< 0.0001 vs the high-diet control group by one-way ANOVA with Dunnett’s post hoc test.

Further detailed posttreatment analysis revealed that Kylo-0603, which was administered at five different doses, significantly reduced the fat content in a dose-dependent manner (Figure 4F). The specific reductions were 8.96%, 9.16%, 14.07%, 26.15%, and, most notably, 44.35%, demonstrating its potent fat-reducing effects (Figure 4G). In addition, Kylo-0603 significantly increased lean mass in mice while reducing fat content. Following an eight-week administration period, there was a notable increase in the percentage of lean mass by 5.32%, 10.95%, 13.52%, 21.94%, and 37.91%, respectively, in comparison to the HFD group (as illustrated in Figure 4H). Given that lean mass is essentially constant (Figure S2B), the observed increase in lean percentage content was not attributable to an increase in lean mass; rather, it was due to a decrease in fat content). These characteristics are not observed with CG-1 administration [40].

### 3.5 Lipid-lowering effects in HFD-induced obese mice

After 10 weeks of Kylo-0603 drug intervention, compared with those in the HFD group, the levels of Chol and LDL-C in the blood of the mice in all the drug-treated groups were significantly decreased in a dose-dependent manner. Eight mice were randomly selected from each group for lipid parameters (including Chol and LDL-C) at week 2 and the end of treatment (week 10). The results of the study demonstrated that Kylo-0603 was effective at reducing plasma cholesterol and LDL-C levels in mice at all the tested doses and that this effect was significantly dose-dependent.

Specifically, after 2 weeks of treatment, mice treated with 3 mg/kg and 10 mg/kg Kylo-0603 presented 52.0% and 64.5% reductions in Chol levels (Figure 5A), respectively, and even more pronounced reductions in LDL-C levels of 84.0% and 85.8% (Figure 5B), respectively, than did the untreated HFD control group. Furthermore, at the end of the 10-week treatment, all the treatments (0.1 mg/kg, 0.3 mg/kg, 1 mg/kg, 3 mg/kg, and 10 mg/kg) significantly inhibited the Chol and LDL-C levels in a dose-dependent manner. Compared with those in the HFD control group, the Chol levels were reduced by 30.6%, 35.9%, 47.9%, 53.9%, and 69.2%, respectively (Figure 5C), whereas the reduction in LDL-C levels was even more significant at 58.0%, 72.7%, 81.8%, 79.7%, and 88.2%, respectively (Figure 5D). At the end of the 2nd and 10th weeks of pharmacological intervention, the remaining 5 mice in each group were subjected to a 6-hour fast, followed by euthanasia. The liver tissues from these mice were collected in isopropyl alcohol for homogenization. Following the intervention with Kylo-0603, alterations in Chol and TG concentrations in the liver were observed. The results demonstrated that Kylo-0603 did not exert a notable influence on the cholesterol content in liver homogenates. However, it was observed to be efficacious in reducing the triglyceride content. In particular, the reduction of triglycerides was most pronounced at a dose of 3 mg/kg, with an average reduction of up to 36.4% (see Figure S3). These data not only reveal the potential of Kylo-0603 to regulate hepatic triglyceride levels under specific conditions, but also strongly demonstrate its potential value in regulating blood lipids and preventing hyperlipidemia and related cardiovascular diseases

**Figure 5.**
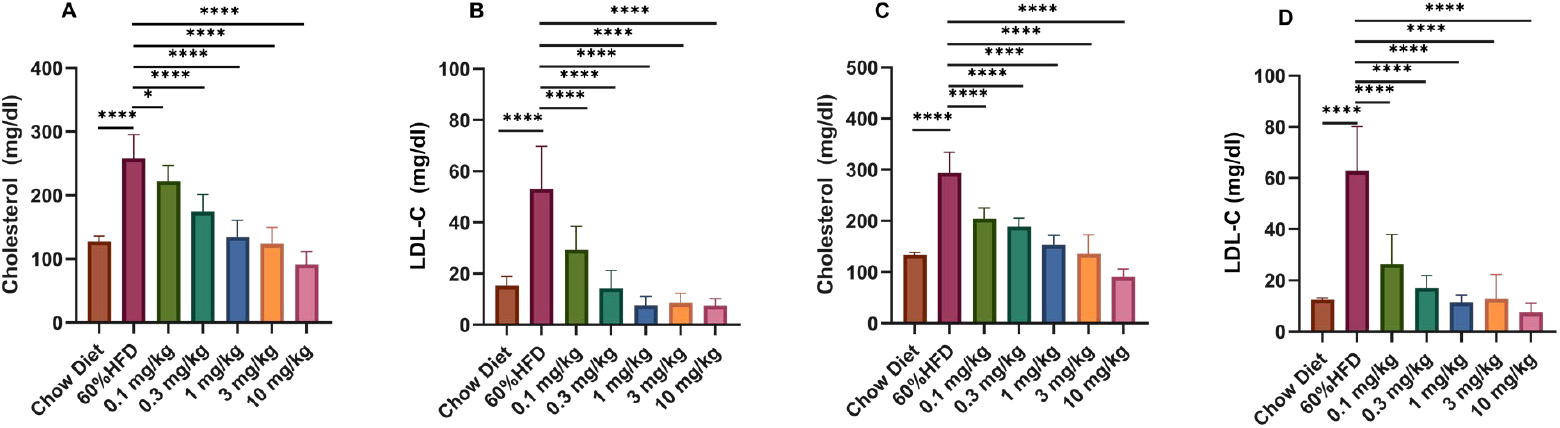
Effects of T3 and GC-1 on blood lipid levels. Cholesterol and LDL-C levels were increased in hypothyroid mice. Treatment of HFD-fed mice with Kylo-0603 decreased cholesterol and LDL-C at all doses. Plasma cholesterol and LDL-C levels in HFD-fed mice after 2 weeks of treatment with Kylo-0603 administration (A and B). Plasma cholesterol and LDL-C levels in HFD-fed mice after 10 weeks of treatment with Kylo-0603 administration (C and D). The data are shown as the means ± SDs; n=5; P< 0.05, **P< 0.01, ***P< 0.001, and ****P< 0.0001 vs the high-diet control group according to one-way ANOVA with Dunnett’s post hoc test.

### 3.6 Efficacy of Kylo-0603 in MASH Mouse Models (F2∼3)

Although the exact molecular pathogenesis of MASH remains to be elucidated, previous studies have shown that CCl_4_ can exacerbate liver fibrosis and damage induced by a high-fat diet (HFD; 60 kcal% fat) [41, 42]. In light of these findings, the present study aimed to explore the potential protective effects of Kylo-0603 on changes in liver structure and function induced by the combination of CCl_4_ and a high-fat diet in 5-week-old male C57BL/6J mice. The experimental design was as follows: Mice were randomized into a control group and an HFD group, both of which received intraperitoneal injections of CCl_4_ at a dose of 0.2 ml/kg twice weekly. After 8 weeks of high-fat feeding, the mice in the HFD group were further divided into six subgroups based on their body weight data at week 8. The results showed that the mice in the CCl_4-_induced control group (HFD+CCl_4_+Vehicle) developed significant hypothyroid-like symptoms, as evidenced by a significant decrease in the blood levels of T3, fT3, and fT4 and an increase in the TSH level. Moreover, the MASH score was as high as 6.3, accompanied by significant inflammation (2.9), ballooning (1.8), and moderate to severe fibrosis (2.6, grades F2-3), meeting the diagnostic criteria for fibrotic MASH. Each dose group (0.1 mg/kg, 0.3 mg/kg, 1 mg/kg, 3 mg/kg, or 10 mg/kg Kylo-0603 in combination with HFD and CCl_4_ solvent received an 8-week treatment. Compared with the control group (HFD+CCl_4_+Vehicle), the Kylo-0603 treatment group presented a significant reduction in the body weight of the mice, and this effect was dose-dependent (reductions ranging from 0.8% to 15.8%). In addition, total cholesterol and LDL-C levels were significantly reduced in a dose-dependent manner (the results of the experiment were similar to those described above for the administration of drugs in HFD-induced obese mice). More importantly, Kylo-0603 effectively reduced the abnormally elevated serum levels of ALT (Figure 6G) and AST (Figure 6H), particularly in the 3 mg/kg dose group, which presented the most significant effects, with ALT and AST decreasing by 73.8% and 76.3%, respectively

**Figure 6.**
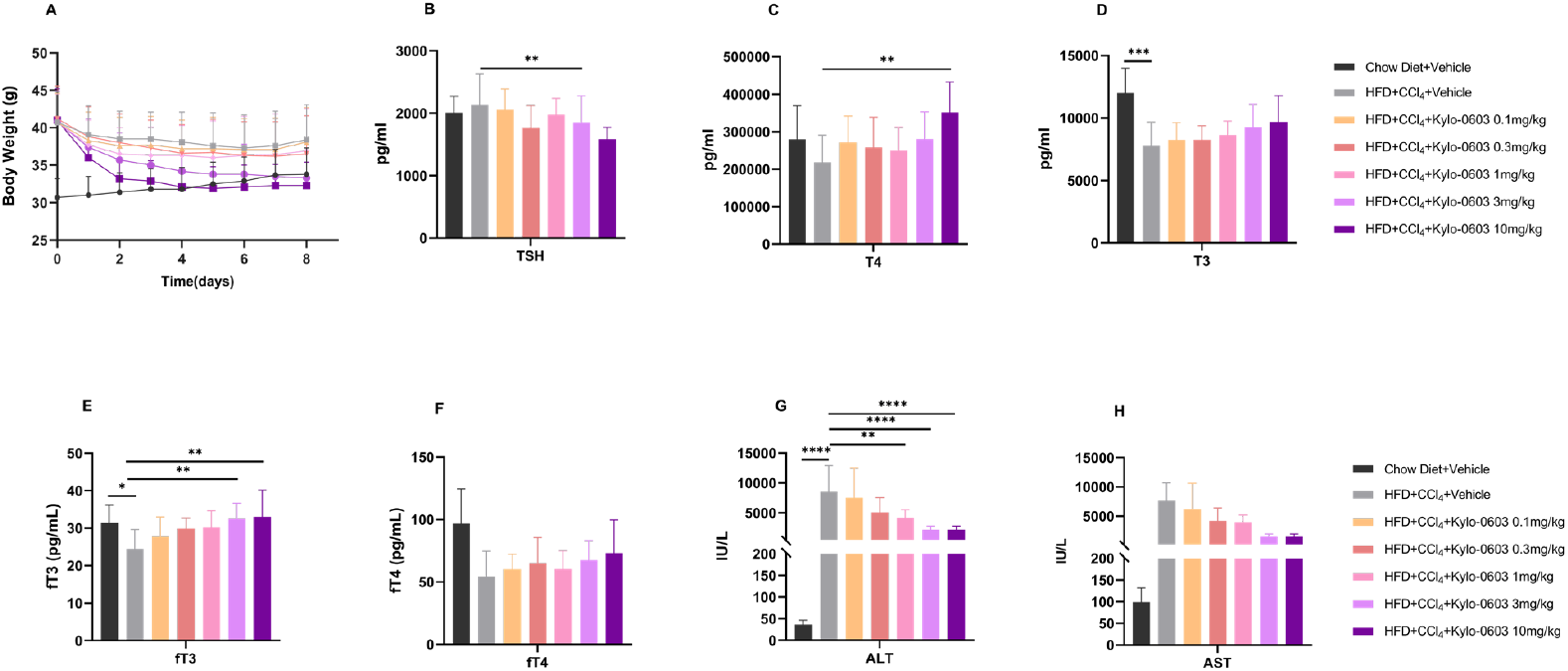
Pharmacological efficacy of Kylo-0603 in MASH mouse models (F2∼3). A shows the body weights of the mice after 8 weeks of drug treatment. B-F show the levels of TSH, T3, T4, fT3, and fT4 in the blood of the mice in each group after 8 weeks of drug treatment. G-H Liver function tests (ALT and AST). The data are presented as the means ± SDs; *, p<0. 05; **, p<0.01; ***, p<0.001; ****, p<0.0001; one-way ANOVA with Dunnett’s post hoc test,n=9.

It is noteworthy that after eight weeks of Kylo-0603 treatment in the MASH mouse models (F2∼3), the levels of thyroid hormones T3, fT3, T4, and fT4 in the drug-treated group of mice exhibited increasing trends to varying degrees, while the TSH levels demonstrated decreasing trends to varying degrees (Figure 6B-F). These findings indicate that Kylo-0603 may facilitate partial improvement in hypothyroidism and mitigate MASH-related pathological alterations in mice. In the HDF-induced obese mouse model, which exhibited no indications of hypothyroidism, the administration of Kylo-0603 did not elicit a notable impact on TSH, T3, T4, fT3, and fT4 levels (data not shown).

The livers from the mice were harvested and evaluated for MASH score and fibrosis using H&E staining, Sirius red staining, and quantitative real-time polymerase chain reaction (qRT-PCR). Following eight weeks of treatment with Ky-lo-0603, the mice in the drug-treated groups exhibited notable improvements in hepatic steatosis, inflammatory response, and ballooning degeneration in comparison to the control group (HFD+CCl_4_+Vehicle), with no exacerbation of hepatic fibrosis. Specifically, at doses of 3 mg/kg and 10 mg/kg, mice exhibited notable reductions in hepatic steatosis and inflammation scores, exceeding 1.0 points (P < 0.001), with decreases in ballooning degeneration scores of 0.7 points (P < 0.05) and 0.8 points (P < 0.01), respectively. Furthermore, the MASH score of the two groups exhibited a notable decline, with a reduction of 3.5 points (P < 0.0001) and 3.7 points (P < 0.0001), respectively (Figure 7A). Additionally, the fibrosis score demonstrated a decrease of 0.6 points (P < 0.05) (Figure 7B). These findings provide compelling evidence that Kylo-0603 not only ameliorates the pathological characteristics of MASH but also effectively inhibits further progression of liver fibrosis. The alterations in hepatic histology were correlated with reductions in plasma cholesterol and LDL-C.

**Figure 7.**
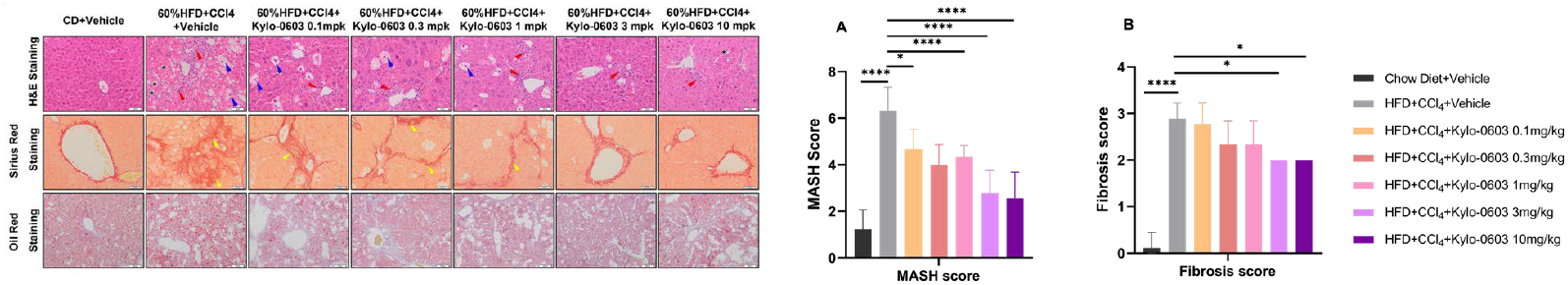
Liver histology of mice after 8 weeks of treatment with either vehicle or Kylo-0603 (0.1–10 mg/kg/day, as indicated). H&E staining was performed. Photomicrographs of representative samples from each treatment group are shown. (*, lipid droplets; blue triangles, balloon-like changes; red triangles, inflammatory foci; yellow triangles, fibrosis) Scale bar=50 μm, n=9. A and B show the MASH scores and fibrosis scores of the liver tissue of the mice after 8 weeks of drug intervention. The data are presented as the means ± SDs; *, p<0. 05; **, p<0.01; ***, p<0.001; ****, p<0.0001; one-way ANOVA with Dunnett’s post hoc test, n=9.

**Table 1.**
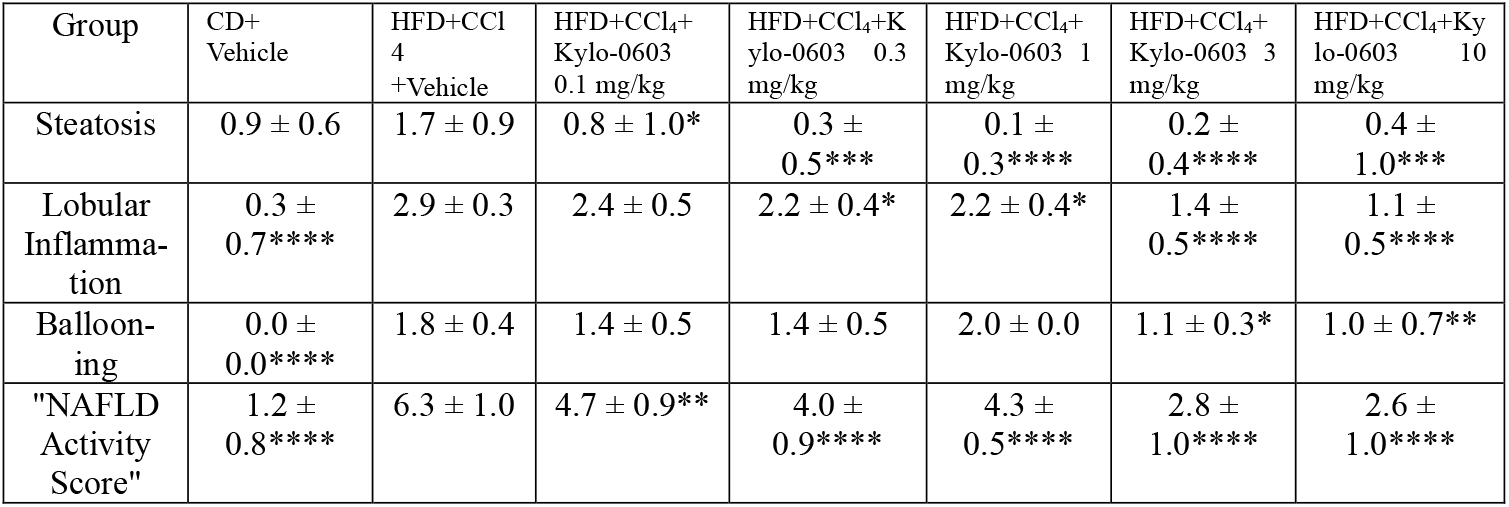
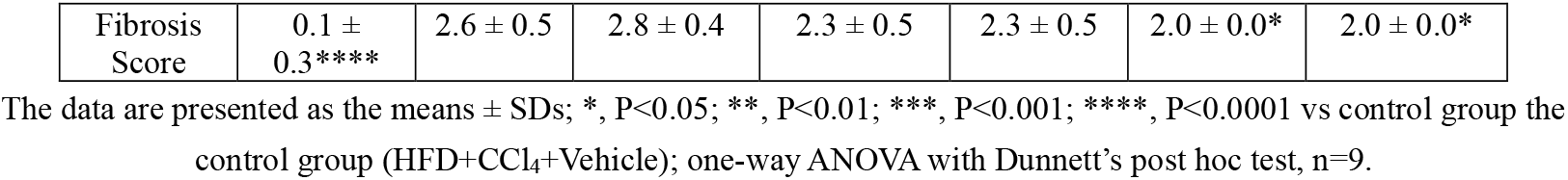
MASH score of liver tissue of mice in each group after 8 weeks of treatment with either vehicle (V) or Kylo-0603 (0.1-10 mg/kg/day)

### 3.7 Accurate assessment of the therapeutic effect of the Kylo-0603 drug via SHG (second harmonic generation)/TPEF (two-photon excitation fluorescence) microscopy technique in MASH mouse models (F2∼3)

As a fundamental component of clinical prognosis in MASH, a precise evaluation of hepatic fibrosis is very crucial. Artificial intelligence-assisted second harmonic generation/two-photon excitation fluorescence (SHG/TPEF) microscopy technique has been introduced because the existing scoring systems are insufficient for capturing the process of hepatic fibrosis regression. This offers a consistent and very precise way to evaluate the pathological characteristics of MASH, especially the serial measurement of collagen fibers and hepatic fibrosis [43]. In this study, we used SHG/TPEF microscopy combined with digital image analysis to evaluate liver tissue sections in a larger scale of tissue area from MASH mouse models (F2∼3) and to analyze the effects of Kylo-0603 on reversing steatosis and inhibiting fibrotic progression in different regions of the liver tissue

By a quantitative analysis of SHG/TPEF images, we assessed in detail the specific effects of different doses of Kylo-0603 (0.1 mg/kg, 0.3 mg/kg, 1 mg/kg, 3 mg/kg, and 10 mg/kg) on fibrosis and steatosis in the MASH mouse models (F2∼3), and the key findings are summarized below:

a. The assessment of fibrosis revealed a notable reduction in the degree of fibrosis in all Kylo-0603-treated groups when compared to the untreated control group (HFD+CCl_4_). Specifically, the percentage of fibrosis decreased from 4.36% in the control group (HFD+CCl_4_) to 2.92% (0.1 mg/kg), 2.12% (0.3 mg/kg), 2.16% (1 mg/kg), 2.03% (3 mg/kg), and 1.25% (10 mg/kg) in the respective dose groups. The improvement between dose groups was statistically significantly different (P < 0.05).
b. Regarding steatosis, Kylo-0603 showed statistically significant improvements between dose groups (P < 0.05) and a substantial reduction in the area of steatosis when compared to the control group at all tested doses. Notably, lipoatrophy was reduced by roughly 60% in the group receiving the lowest dose (0.1 mg/kg), and by a considerable 90% in the group receiving the highest dose (10 mg/kg). These results show how effective Kylo-0603 is at preventing lipoatrophy. 21.66% (HFD+CCl_4_), 5.07% (0.1 mg/kg), 4.39% (0.3 mg/kg), 4.19% (1 mg/kg), 2.22% (3 mg/kg), and 1.86% (10 mg/kg) were the percentages of lipoatrophy, respectively.

### 3.8 Gene expression analysis in the liver of the MASH mouse models (F2∼3)

To explore the mechanism by which Kylo-0603 promotes lipid metabolism, we used Illumina’s NEBNext® UtraTM RNA Library Prep Kit to examine the expression of relevant genes in the livers of the MASH mouse models (F2∼3) after Kylo-0603 administration (Figure 8). We first examined the expression of THR-regulated genes, including iodothyronine deiodinase 1 (Dio1), malic enzyme 1 (Me1), and thyroid hormone responsive (Thrsp). Me1 encodes an enzyme that catalyzes the conversion of malate to pyruvate while concomitantly generating NADPH from NADP, and its upregulation promotes energy metabolism [44]. Thrsp is mainly a nuclear protein induced by thyroid hormones, carbohydrate intake, adipose tissue differentiation and lactation and plays an important role in the regulation of lipid metabolism. The Dio1 gene in liver tissue is responsible for the conversion of T4 to T3, thereby facilitating intrahepatic energy metabolism. Reduced DIO1 levels and activity have been observed in humans and rodents with advanced MASH, and DIO1 knockdown results in increased hepatic lipid content, suggesting that downregulation of DIO1 may exacerbate hepatic lipid accumulation and MASH progression. Kylo-0603 significantly upregulated the expression of Dio1, Thrsp and Me1 in a dose-dependent manner in the MASH mouse models (F2∼3) (Figure 9A). LDL-R (low-density lipoprotein receptor) promotes the cellular uptake of LDL and facilitates cholesterol degradation. Kylo-0603 upregulated LDL-R gene expression in a dose-dependent manner. This finding was consistent with a dose-dependent reduction in the serum LDL-C level (Figure 9A). Kylo-0603 significantly downregulated the expression levels of genes closely related to inflammatory factors in liver tissues, which in turn effectively reversed the inflammatory response in liver tissues and significantly improved the pathology of MASH syndrome. This therapeutic effect was validated by assessing liver histopathology in animal models. Kylo-0603 inhibited the genes encoding the inflammatory factors Tnfrsf1a, IL6, IL17rb, and IL17ra (Figure 9B). We thoroughly analyzed the dynamics of gene expression in liver tissues closely related to fibrosis, focusing on collagen synthesis genes that play a central role in the pathological process of fibrosis. The results revealed that the expression of these genes was significantly suppressed with increasing doses of the Kylo-0603 drug, demonstrating a distinct dosedependent characteristic (Figure 9C). These findings directly demonstrate that Kylo-0603 can effectively reduce abnormal collagen deposition in the liver by regulating collagen synthesis. This mechanism not only provides intuitive evidence for the active role of the drug in the treatment of liver fibrosis but also strongly supports its potential to alleviate or even stop the fibrotic process.

**Figure 8.**
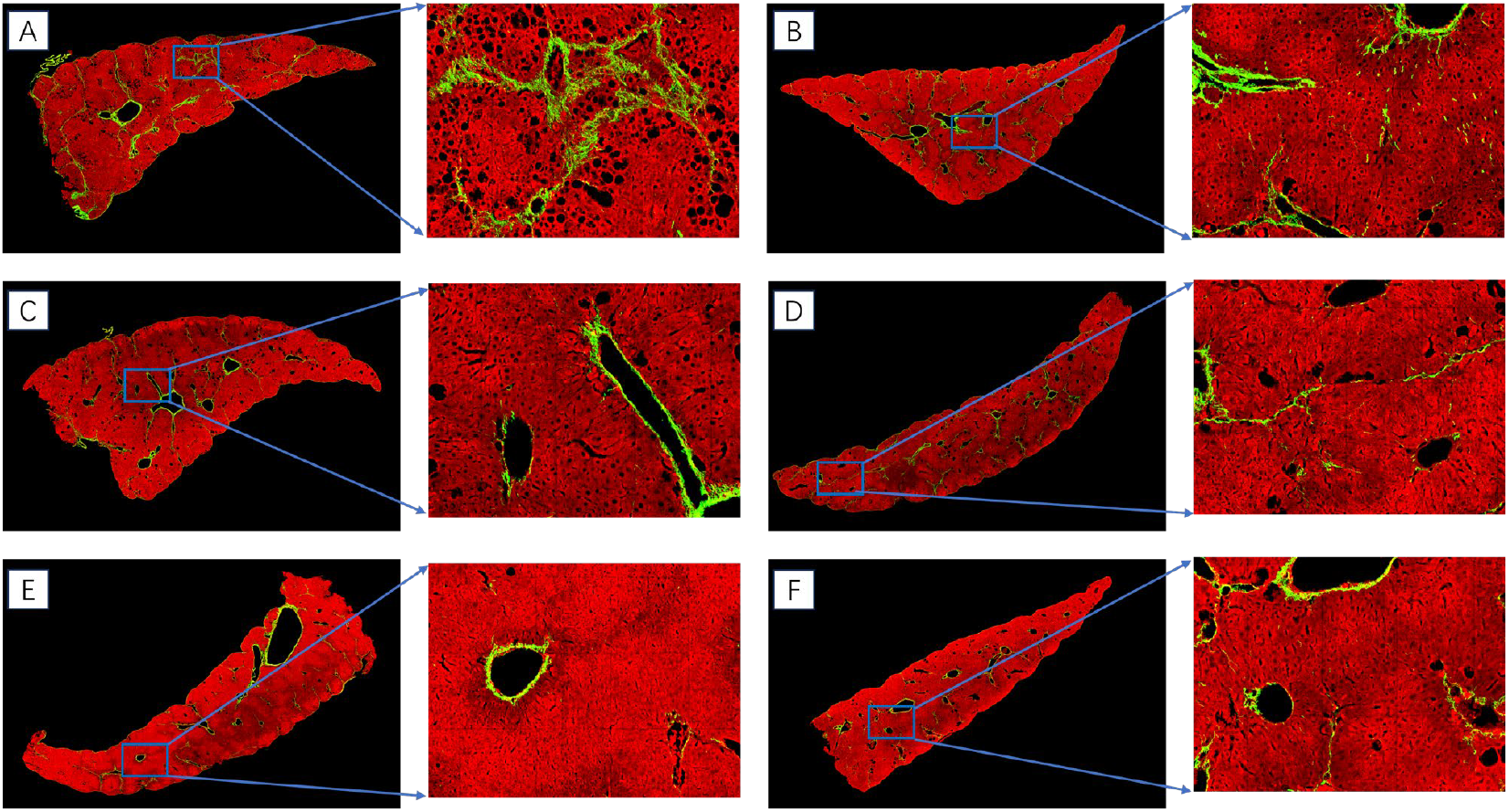
Effect of Kylo-0603 drug on the reversal of steatosis and inhibition of fibrotic process in liver tissues of the MASH mouse models (F2∼3), analyzed by SHG/TPEF microscopy combined with digital image analysis technique. A shows the control and untreated groups (SHG fibrosis (green): 4.36%, steatosis (black lipid cavity and surrounding involved area):21.66%); B shows the sample image after 0.1 mg/kg administration (SHG fibrosis (green): 2.92%, steatosis (black lipid cavity and surrounding involved area):5. 07%); C shows the sample image after 0.3 mg/kg administration (SHG fibrosis (green): 2.12%, steatosis (black lipid cavity and surrounding involved area): 4.39%; D shows the sample after 1 mg/kg administration (SHG fibrosis (green): 2.16%, steatosis (black fat space and surrounding area): 4.19%); E shows the specimen after 3 mg/kg administration (SHG fibrosis (green): 2.03%, steatosis (black fat vacuoles and surrounding areas): 2.22%; F shows the sample after 10 mg/kg administration (SHG fibrosis (green): 1.86%, steatosis (black fat vacuoles and surrounding areas): 1.25%

**Figure 9.**
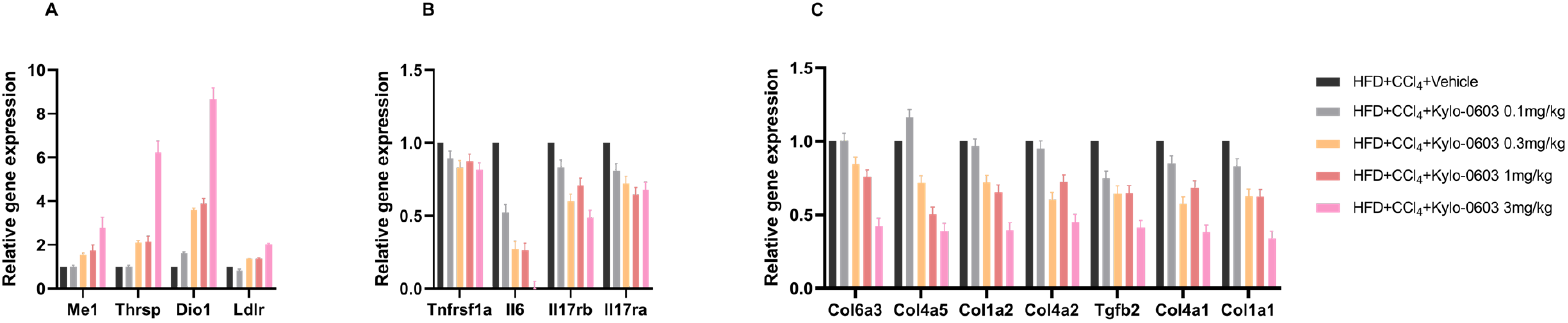
Effects of Kylo-0603 on the expression of THR-related genes, lipid metabolism genes, inflammatory factor genes, and fibrosis-related genes in the liver of the MASH mouse models (F2∼3). Total RNA was extracted from the frozen liver tissue of the MASH mouse models (F2∼3), and the mice were injected daily with saline or Kylo-0603 at a dosage of 0.1–3 mg/kg/day. Transcriptome sequencing is based on the Illumina sequencing platform. The relative expression of a gene was calculated by normalizing the true expression level of the target gene to the expression level of that gene in the control group. A shows the trend of metabolism-related gene expression changes in liver tissue. B shows the inhibitory effect of Kylo-0603 on the inflammatory factor genes Tnfrsf1a, Tnfrsf11a, IL6, IL1b, IL17rb, IL17ra and Cxcl2. C shows the expression of collagen synthesis-related genes in liver tissue.

Notably, Kylo-0603 stimulated the upregulation of peroxisome-related genes in a dose-dependent manner. These results demonstrated the unique ability of Kylo-0603 to improve MASH symptoms, which may be largely attributed to its unique liver-targeting properties.

## 4. Discussion

MASLD is currently the most common liver disease worldwide and a major cause of liver-related morbidity and mortality. According to statistics, the prevalence of MASLD has reached as high as 38% [45], and it is still rising at a startling rate. Because of its intricate pathophysiology, MASH, a severe manifestation of MASLD, presents substantial obstacles to the advancement of clinical studies and the creation of new medications. Currently, only one MASH drug, Resmetirom, was approved for marketing in the United States on March 14th this year, primarily for the treatment of adults with noncirrhotic MASH with moderate to advanced liver fibrosis (consistent with stages F2 to F3 fibrosis) in the USA [46]. In vitro THR competition assays demonstrated that Resmetirom exhibited the highest degree of THR-β selectivity among thyromimetic agents, with a selectivity ratio of 12 or 17 for THR-β over THR-α. GC-1 exhibited a selectivity ratio of approximately 0.9 to 10 for THR-β over THR-α, while VK2908A (MB07344) demonstrated a selectivity ratio of 2.5 to 15.8 for THR-β over THR-α. These findings were previously published [28,47-48, 54]. In this work, we developed the drug Kylo-0603, which, although not optimal in terms of THRβ selectivity, is endowed with unprecedented liver-targeting properties owing to its unique GalNAc structural design [38, 49]. This innovative design ensures that the drug can be efficiently and specifically transported to the liver, significantly reducing potential side effects on other nontarget tissues in the body, with drug concentrations in the liver measured to be nearly 70,000 times higher than those in the blood at 0.25 h after a single dose in Rats. For example, the tissue/plasma ratios of GC-1 and T3 in the liver are similar [40]. Owing to its unique high hepatic targeting ability, Kylo-0603 shows exceptional properties in treating MASH disease.

Obesity is an important medical challenge that not only affects the quality of life of individuals but also significantly increases morbidity and mortality from many diseases. Current treatments for obesity are effective but still have significant limitations in terms of safety and efficacy, which has prompted researchers to explore new therapeutic strategies continuously. Among them, thyroid hormone receptor modulators have attracted much attention for their potential therapeutic effects. Thyroid hormones exert their physiological effects through two main receptors, TRα and TRβ. TRα mainly mediates the effects of thyroid hormones on the heart, whereas TRβ is strongly associated with the control of obesity and hypercholesterolemia, and its activation does not trigger the undesirable side effects of thyroid hormones on the heart, bones, or skeletal muscles. However, the action of endogenous thyroid hormones is nonselective and can lead to several adverse effects, including cardiac stimulation.

In this study, Kylo-0603, a novel drug, demonstrated significant potential for weight loss and lipid reduction. The drug effectively reduced the body weight of high-fat diet-induced obese mice, especially at doses of 3 mg/kg and 10 mg/kg. Moreover, Kylo-0603 also significantly reduced the amount of fat in the mice, a mechanism similar to that previously studied for T3 and GC-1 weight reduction through increased oxygen consumption. However, Kylo-0603 is unique because of its ability to target the liver. This means that the drug acts primarily on TRβ in the liver, avoiding the adverse effects that nonselective thyroid hormones can cause in organs such as the heart. As a result, Kylo-0603 has a more pronounced advantage in weight loss than GC-1. In addition, Kylo-0603 has unique advantages over the currently popular GLP-1-targeting drugs. Although GLP-1 drugs can control blood sugar and reduce body weight to a certain extent, their mechanism of action is relatively complex, and 40% of their body weight reduction is muscle volume [50,51]. In contrast, Kylo-0603 achieves more precise and effective weight loss and fat reduction effects by acting directly on THRβ in the liver and reducing only fat mass. Thyroid status strongly influences contractile function and the mass of skeletal muscle. However, Kylo-0603 has high liver-specific targeting ability and selectivity for THRβ, and it has been demonstrated to not cause T3-mediated fiber type shifts or nutrient changes related to bone mineral density (BMD/BMC) ((Figure S2). In a high-fat diet-induced obese mouse model, we evaluated the effects of Kylo-0603 on lipid metabolism. The results showed that after 2 to 10 weeks of Kylo-0603 intervention, Kylo-0603 was able to significantly reduce blood Chol and LDL-C levels in mice compared with those in HFD controls and that this reduction was dose-dependent. This finding is consistent with previous results with THR-β-selective agonists such as GC-1 and MGL-3196, both of which have similar effects. Moreover, the lipid changes induced by Kylo-0603 were consistent with the changes in the expression of genes related to hepatic lipid metabolism (Figure 6). In particular, similar to previous studies showing that both T3 and GC-1 significantly increased the expression of Dio1 and Me1, the Kylo-0603 study yielded consistent results. In addition, Kylo-0603 upregulated the expression of Thrsp. These results suggest that Kylo-0603 may act through a THR-dependent signaling pathway as T3. LDL-R is crucial to the metabolism of LDL particles. LDL-C combines with LDL-R on the surface of the liver cell membrane and enters human liver cells through endocytosis. In mice with high-fat diet (HFD)-induced obesity models or MASH mouse models (F2∼3), Ky-lo-0603 demonstrated a significant and dose-dependent reduction in plasma cholesterol and LDL-C levels. This finding aligns with the mechanism by which thyroid hormones and thyroid hormone analogs reduce cholesterol levels by increasing the expression of hepatic LDL-R [48,52].

In addition to the aforementioned studies on body weight and lipids, we also investigated the impact of the Ky-lo-0603 drug on liver fibrosis. The results demonstrated that after 8 weeks of treatment with Kylo-0603, H&E staining revealed significant improvements in fat content, inflammation, and ballooning in liver tissues, indicating that Kylo-0603 improved MASH symptoms without worsening fibrosis. To gain further insight into the pharmacodynamic properties of Kylo-0603 in alleviating MASH symptoms, we employed advanced TPE/SHG (two-photon excitation/second harmonic generation) imaging technology to provide an in-depth analysis of improvements in liver fibrosis. The analysis demonstrated that the compound designated Kylo-0603 exhibited notable therapeutic efficacy at specific doses in the MASH mouse models. The analysis demonstrated that the compound designated Kylo-0603 exhibited significant therapeutic efficacy at specific doses in the MASH mouse models. The drug was effective in reversing steatosis and also showed the ability to halt the progression of fibrosis, thereby highlighting its potential for therapeutic applications in various fields.

This paper presents a comprehensive examination of the expression of genes associated with lipid metabolism and fibrosis, along with an extensive analysis of the expression changes of numerous related genes. For instance, impaired peroxisome function is linked to the development of MASH [53]. This study reveals that Kylo-0603 upregulates peroxisome-related genes in a dose-dependent manner. This finding further confirms the unique effect of Kylo-0603 in alleviating MASH symptoms, primarily due to its distinctive hepatic targeting properties. For additional details regarding the expression of other relevant liver genes, please refer to the heatmap in the Supporting Material.

Notably, T3-like binding to THR-α of THR-β agonist has been demonstrated to exert deleterious effects on the skeletal and cardiac systems, which represents a significant impediment to its utilization in the treatment of obesity and MASH. The novel THR-β agonist Kylo-0603, developed in this study, can specifically target the liver, thereby effectively circumventing the potential adverse effects that may be induced by nonspecific binding of THRα in cardiac tissues. In the MASH mouse model induced, treatment with Kylo-0603 resulted in significant improvement in hypothyroidism and a notable reduction in abnormally elevated levels of alanine aminotransferase (ALT) and alanine aminotransferase (AST) (Figure 6). Furthermore, this effect was dose-dependent. In conclusion, these data demonstrate that Kylo-0603 is an effective agent for reducing the degree of hepatic injury and improving hypothyroidism induced by CCl_4_ injection.

In conclusion, this study designed and synthesized an innovative liver-targeting compound, Kylo-0603, with unique both liver-targeting and THR-β agonist properties. Kylo-0603 was crafted through precise chemical design, connecting three galactosamine structures to a T3-like structure via specific ester or amide bonds, resulting in an efficient liver-targeting molecule. Experimental validation demonstrated that Kylo-0603 exhibits exceptional stability in plasma, high selectivity for THRβ, and excellent pharmacokinetic and safety characteristics. This research breakthrough not only opens up a new avenue for exploring therapeutic drugs for MASLD and MASH but also lays a solid scientific foundation for potentially significant breakthroughs in improving the quality of life for patients with liver diseases in the future.

## Author contributions

Kunyuan Cui, serving as the corresponding author of this research. Xueqin Lu primarily takes charge of chemical compound design, Pharmacology, PK and safety studies. Shengjun Wang and Yanchun Du devoted themselves to the chemical synthesis of the compound presented in this article. Bixian Xie mainly engages in analytical chemistry research. Qingyan Chen specializes in the field of Pharmacology. Jinzhen Lin serves as the chemistry advisor. Bailing Chen completed the data collection, organization and writing of the article.

## Acknowledgments

This work was supported by Hygieia (Hangzhou) Pharmaceuticals and Kylonova (XiaMen) Biopharma Pharmaceuticals.

